# Self-organization in natural swarms of *Photinus carolinus* synchronous fireflies

**DOI:** 10.1101/2021.01.26.428319

**Authors:** Raphaël Sarfati, Julie C. Hayes, Orit Peleg

## Abstract

Fireflies flashing in unison is a mesmerizing manifestation of animal collective behavior and an archetype of biological synchrony. To elucidate synchronization mechanisms and inform theoretical models, we recorded the collective display of thousands of *Photinus carolinus* fireflies in natural swarms, and provide the first spatiotemporal description of the onset of synchronization. At low firefly density, flashes appear uncorrelated. At high density, the swarm produces synchronous flashes within periodic bursts. Using three-dimensional reconstruction, we demonstrate that flash bursts nucleate and propagate across the swarm in a relay-like process. Our results suggest that fireflies interact locally through a dynamic network of visual connections defined by visual occlusion from terrain and vegetation. This model illuminates the importance of the environment in shaping self-organization and collective behavior.

---

The spontaneous synchronization of thousands of flashing fireflies is a natural spectacle that elicits fascination, even bewilderment [1]. Early scientists investigating popular accounts of firefly synchrony often dismissed it as an illusion, a statistical accident, or an observational artifact, such as the observer’s blinking eyelids or the sudden alignment of the fireflies’ lanterns (light-producing organs) from the wind [2]. Skepticism might have persisted for a few decades because these displays are quite rare, and as a natural occurrence, synchronous patterns can be complex and noisy. But careful studies over the past 50 years, facilitated by new imaging techniques and analytical tools, have confirmed that precise synchrony does occur in swarms of specific species under proper circumstances [3, 4, 5, 6].

Fireflies use flashes for species recognition and courtship [7]. Typically, males advertise their fitness by flying and flashing, while females respond selectively from the ground [8]. In a few species, and generally associated with a high swarming density, males tend to synchronize their rhythmic flashing with their peers. Synchronous flashing is a compelling display of collective behavior, and a readily-accessible example to study synchrony in natural systems. This is why firefly synchrony has often been cited as an inspiration for the theoretical study of systems of coupled oscillators, such as the Integrate-and-Fire, Winfree, or Kuramoto models [9, 10], which have generated an abundant literature [11]. However, even though synchronous fireflies are directly observable, the connection between theory and natural patterns has rarely been attempted rigorously [12]. In fact, spatiotemporal data currently available shows that these models in their current form are unable to explain a wide variety of natural features of firefly synchrony [5, 6].

To reconcile theory with empirical observations, we video-recorded the collective flashing display of *Photinus carolinus* fireflies in Great Smoky Mountain National Park during peak mating season in June 2020. The fireflies’ primary natural habitat are the densely-forested creeks of the Elkmont, TN area of the park. We positioned our cameras at the edge of a small forest clearing, facing a steep ridge (Fig. 1A). Using stereoscopic recordings, flash occurrences were localized in three-dimensional space (Fig. 1B). After camera calibration and flash triangulation [13], we were able to reconstruct, for each night, a cone-shaped portion of the swarm (30m long and up to 10m wide) containing up to half a million space-time coordinates (Fig. 1C). It appears that flashes tend to correlate strongly with terrain geometry, indicating that fireflies localize primarily in a thin layer about 1m above ground (Fig. 1D), in agreement with our previous observations [6]. This layer is crowded with bushes and short vegetation. Therefore, this camera placement provides an external, global view of the swarm that is quite different from the perspective of a single swarming firefly. As the natural swarm extends over hundreds of meters, and visual occlusion from vegetation is significant, these reconstructions also constitute only partial renderings of the swarm.

**Fig 1.**
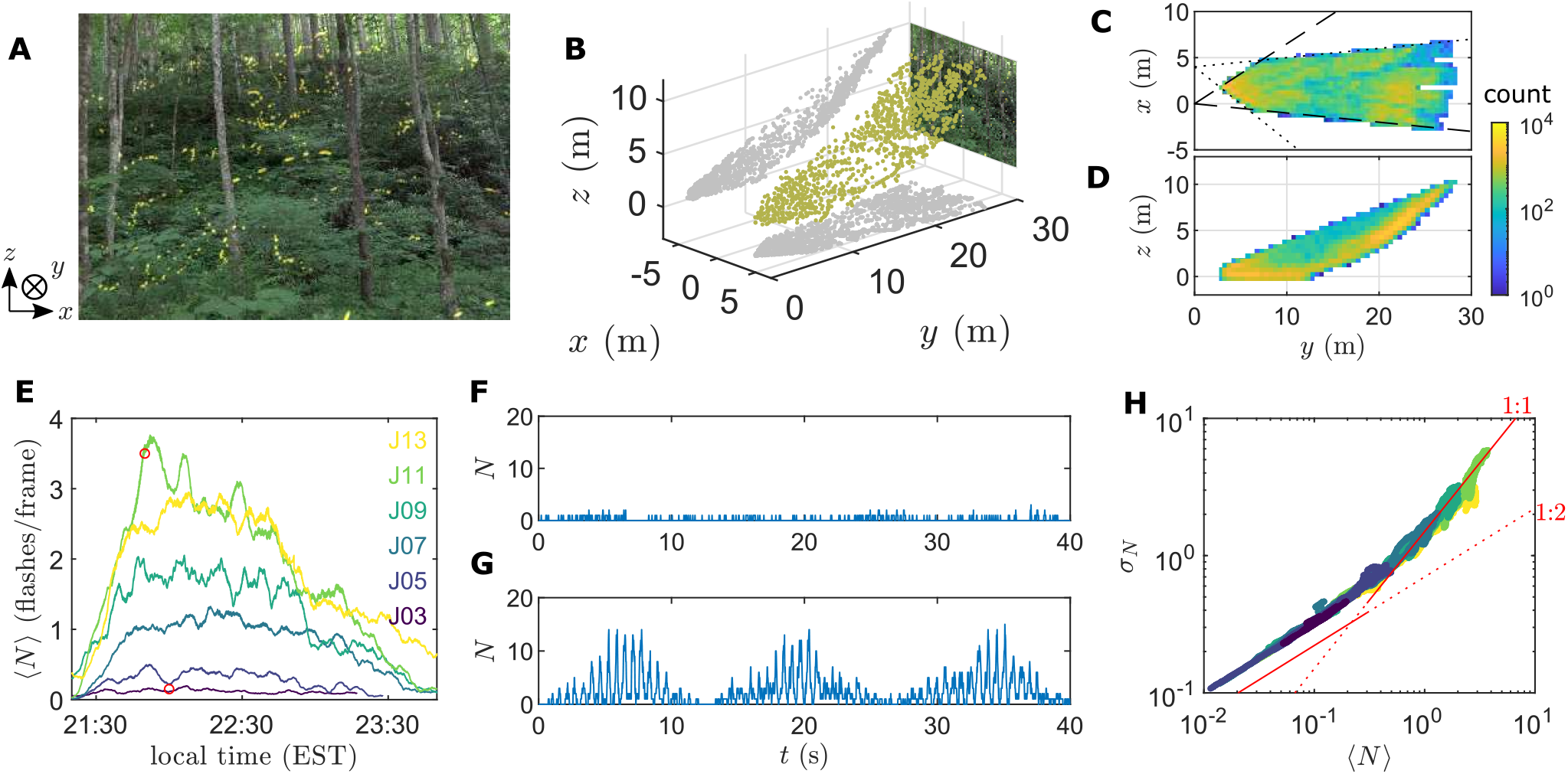
Three-dimensional reconstruction of a natural swarm and density-dependent collective flashing. **(A)** View from one camera, showing some *Photinus carolinus* flashes in their natural habitat (composite image for illustration). A steep ridge covered by dense vegetation is visible in the background. **(B)** Using a second camera to record the same scene (60 frames-per-second), flash occurrences can be located in 3D (yellow dots; 2D projections in grey). The *x*− and *y*−axes define the horizontal plane, with *y* pointing away from the cameras and towards the ridge; the *z*−axis is vertical upwards. **(C, D)** Spatial distribution of 3D-reconstructed flash occurrences. Colors indicate number of flashes within (0.5m)^2^ bins, on a logarithmic scale. Projection in the horizontal plane shows the swarm from above (C). Due to the cameras’ limited fields-of-view (dashed and dotted lines), only a cone-like portion of swarm can be reconstructed. Projection in a vertical plane perpendicular to the ridge (D) shows that fireflies localize strongly in a 1m layer above ground, including along the slope of the ridge. **(E)** Moving averages (over 5min) of the number of flashes per frame, ⟨*N* ⟩, for each night of recording (June 3-13). For clarity, only odd nights are shown. Firefly density increases steadily every day, until peak is reached (June 10 to 13). **(F, G)** Time series of *N* for a short interval around 21:45 (red circles in E), for a low-density night (F, June 3) and high-density night (G, June 11). At low ⟨*N* ⟩, flashes occur uniformly with little variation. At high ⟨*N* ⟩, *N* exhibit large fluctuations, with flash occurrences clustering at specific times. Fireflies flash synchronously every ∼0.5s during periodic bursts repeated every ∼12s. **(H)** Scaling of the standard deviation *σ*_*N*_ with the mean ⟨*N* ⟩, for all nights displayed in (E). All available data collapses on a single curve. At small ⟨*N* ⟩, *σ*_*N*_ ∼ ⟨*N* ⟩^1*/*2^. At large ⟨*N* ⟩, *σ*_*N*_ ∼ ⟨*N* ⟩. Red lines of slope 1/2 and 1 are shown as a guide.

*P. carolinus* fireflies produce, individually, flashes of 10-15ms repeated up to 8 times, while either immobile or in flight [5, 6]. They are active for approximately 3 hours every night after sunset during about 2 weeks in early summer [8]. We recorded from June 3, when a few rare flashes started to be seen, to June 13, the 4^th^ consecutive night of peak activity, as evidenced by the averaged number of flashes *N* recorded in a given frame (Fig. 1E). Synchronous flashing appears to necessitate a critical density of fireflies to occur. When only few fireflies are active (June 3-5, and early or late in the night), collective flashing appears incoherent (Fig. 1F). During peak nights, flashes tend to cluster at specific times, as *N* exhibits a doubly-periodic pattern of synchronous flashes every 0.55s during repeated bursts lasting about 10s (Fig. 1G) [14]. These two features have well-defined frequencies evidenced in their corresponding power spectra (Fig. S1). However, while Fourier transforms reveal periodicity, they do not inform about synchrony. To quantify the onset of synchrony, we study the distribution of *N* over 5min-time intervals (Fig. 1H). At low density, the standard deviation of *N, σ*_*N*_, scales sublinearly with the mean ⟨*N* ⟩, and approximately as 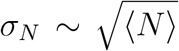, suggesting that flashes are randomly distributed. Past a certain density threshold, however, the scaling becomes linear, *σ*_*N*_ ∼ ⟨*N* ⟩, indicating clustering [15]. This marks synchronous behavior.

This density-dependent transition from disorder to order has been observed in various animal systems exhibiting collective behavior and is a feature of common mathematical models [16]. The underlying mechanism is often believed to stem from an increasing pressure to follow the behavior of the peers when they are numerous. However, the fireflies’ cooperative behavior at high density does not seem to be the result of increasing peer-pressure, since even two fireflies in isolation attempt to flash in synchrony, as previously demonstrated in controlled experiments [6]. Rather, we hypothesize that fireflies interact at short range, therefore high density is necessary to receive and relay the flash information across the swarm.

To explore this hypothesis, we investigate the common observation that the collective flashing of *P. carolinus* appears to be “propagating” or “cascading” across the terrain [17]. Specifically, as flash bursts start and end with only a few flashers (Fig. 1G and Ref. [6]), it has often been reported, but never measured, that bursts tend to originate and terminate at distinct locations many meters apart, often the top and bottom of a ridge. Flashes are sometimes perceived to move progressively from one location to the other over the course of a burst, then repeat the same path during a few minutes. Hence, we look for evidence of such flash propagation in our 3D data. Since the signal is noisy but periodic, we first calculate each flash’s relative timing within a burst. For each burst, we define the time with the maximum *N* as the origin [13]. Each flash occurrence within this burst is then labelled by a phase *ϕ*, roughly between -5s and +5s, corresponding to its relative time in the burst (Fig. 2A). By doing so, we were able to identify certain time intervals of a few minutes that show a clear propagation up and down the ridge over the course of ≃10s, when averaged over ∼ 50 bursts (Movie S1 and S2). In Fig. 2B, for example, early flashes are concentrated at the bottom of the ridge, and late ones at the top. The distribution of *ϕ* along the direction perpendicular to the ridge (*y*-axis) follows a linear progression (Fig. 2C).

**Fig 2.**
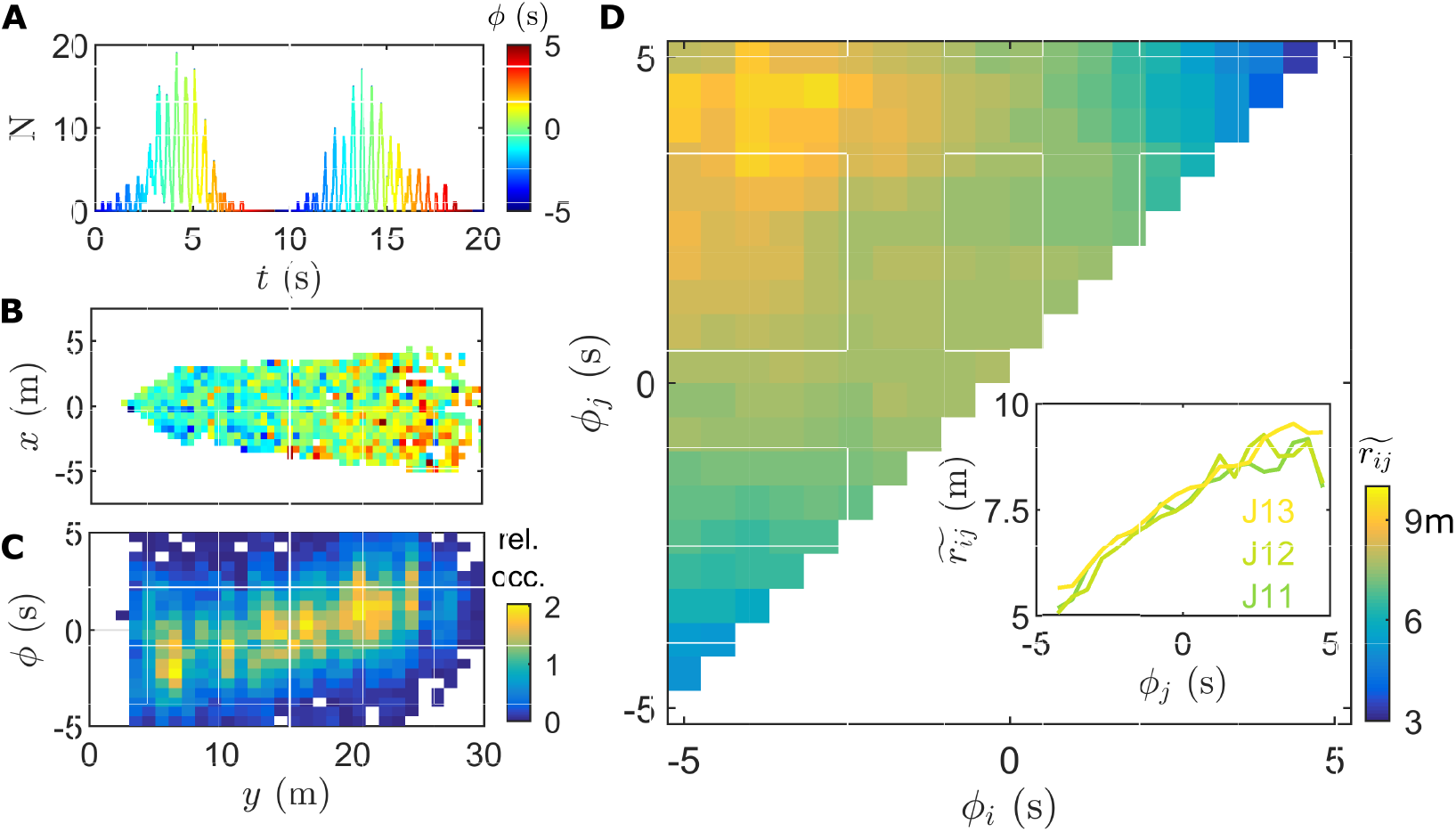
Flash propagation across the swarm. **(A)** Flash occurrences are associated with a phase *φ* indicating their relative timing within a burst. The center of the burst (highest peak) is defined as *ϕ* = 0s. **(B**,**C)** Example of flash propagation along the *y*-axis over a 10min interval (11pm June 10). On average, early flashes are located at the bottom of the ridge (close to the cameras) while late flashes are located at the top. (B) Average *ϕ* in 0.5×0.5m^2^ space bins, same colors as (A). Bins close to the cameras show a negative phase, while bins far have a positive phase. (C) Distribution of *ϕ* along the *y*-axis. Colors indicate relative occurrence. **(D)** Flash propagation over 150min, June 11. For each pair of intraburst flashes occurring at (*ϕ*_*i*_, *ϕ*_*j*_) corresponds a distance *r*_*ij*_. In each bin of the (*ϕ*_*i*_, *ϕ*_*j*_) matrix, the distributions of distances are represented by their median value, 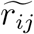, ranging from 3m to 10m. The diagonal has been removed to avoid intraflash self-correlations. (Inset) Median distance 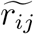 as a function of *ϕ*_*j*_ for *ϕ*_*i*_ = 5 ± 0.25s (leftmost column in the larger plot), for June 11-13. The increase in 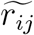 by about 5m over 10s is approximately linear in time.

However, flash propagation does not always follow a specific terrain orientation. In order to characterize flash propagation in general and without relying on a specific coordinate system, we calculate the distribution of distances *r*_*ij*_ between pairs of *intraburst* flashes occurring at *ϕ*_*i*_ and *ϕ*_*j*_ *> ϕ*_*i*_. When all flash occurrences from a night’s recording are processed together [13], a pattern emerges: the median distance 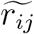 between flashes increases linearly over time, especially relative to early flashes (Fig. 2D). This defines a propagation velocity for the burst activation front. The small value of 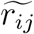 at *ϕ*_*i*_, *ϕ*_*j*_ ≃ −5s and *ϕ*_*i*_, *ϕ*_*j*_ ≃ +5s indicates that very early and very late flashes are strongly localized. The same trend is observed for all data corresponding to peak nights (Fig. 2D Inset). This analysis suggests that flash propagation occurs *on average* at constant speed of about 0.5m/s. Importantly, flash propagation is not associated with any significant flow of flashing fireflies along the propagation path (Fig. S2). This demonstrates that flash propagation occurs through a relay-like process, similar to a wave, whereby information, not matter, is transported.

Therefore, burst propagation suggests that active fireflies interact with the swarm locally, rather than globally. This is common and well-documented in animal groups. Local interactions have been proposed to be either metric [18], where individuals interact with peers within a certain distance, or topological [19], where interactions occur through a set number of nearest neighbors. In flocks, schools, and crowds, local interactions have been shown to result in a constant-velocity (linear), underdamped propagation of information [20, 21, 22, 23].

What is remarkable within *P. carolinus* swarms is that information propagation is linear only *on average*. In fact, simultaneous flashes, even early and late ones, can be spatially distant. In other words, the distributions of *r*_*ij*_(*ϕ*_*i*_, *ϕ*_*j*_) are broad (Fig. S3), in stark contrast to bird flocks or human crowds where the time-distance relationship is very narrow [20, 21, 22, 23]. This suggests that local interactions may extend beyond each firefly’s immediate geometric vicinity.

As further evidence, we find that, although collective flashing is symmetric within a burst (Fig. 3A), firefly movement is not. We consider streaks, defined as the spatial path of a flash as it appears on successive frames (typically 5 to 8 frames). We find that early streaks (*ϕ <* 0) move significantly faster than late ones (*ϕ >* 0), as seen on Fig. 3B. This is similar to what had been observed previously in controlled experiments within an unobstructed confining volume, where the burst leader was flashing longer and flying farther than followers [6]. Most importantly, motion asymmetry suggests that at least some fireflies are able to perceive the global state of the swarm (*i*.*e*. relative delay in the burst), and not simply their local environment. Otherwise, purely local sensing would not create collective behavior disparities during bursts.

**Fig 3.**
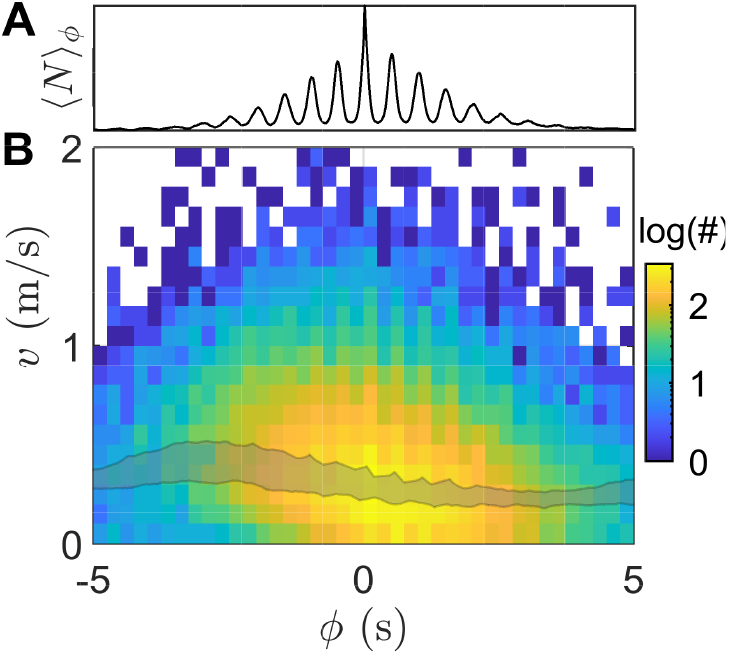
Firefly movement during bursts. **(A)** Average burst, obtained by averaging *N* over the *ϕ*-space, ⟨*N* ⟩ _*ϕ*_. Flash distribution is almost perfectly symmetric. **(B)** Distribution of streak velocities as a function of streak phase *ϕ*, June 10. Colors indicate count, in log10 scale. Early streaks (*ϕ <* 0) are significantly faster than late ones (*ϕ >* 0). Median values for peak nights (June 10-13) all fall within the shaded area.

This leads us to hypothesize that local interactions may have a complex structure, and notably involve a wide range of distances. To explain this framework, we consider the line-of-sight premise: fireflies can only interact with peers that are directly in their field-of-view, *i*.*e* when a pair is connected by an unobstructed line. In dense groups of large animals, visual occlusion is created by nearest neighbors, which are generally concentrated at a characteristic distance. Therefore, local interactions, whether metric or topological, must have a narrow distribution of distances. In dilute swarms, however, there is no such “screening”, and lines-of-sight may be widely distributed. For example, pair interactions in mosquito swarms typically do not follow a characteristic scaling [24].

*P. carolinus* congregations present an intermediate situation: the swarm is dilute, but the environment creates significant visual occlusion. Elevated fireflies have access to a wider view of the swarm, but they are few. The majority of fireflies are located among the vegetation, and may have their field-of-view significantly obstructed. To elucidate the type of interactions that fireflies can establish, we used local swarm reconstructions obtained from 360-degree cameras placed directly within the vegetation, 0.6m above ground [13]. This setup offers an immersive view from the perspective of an “average” swarming firefly. In particular, it allows to characterize the distribution and range of line-of-sight interactions (within a scaling factor relating the light sensitivity of the cameras and of the firefly’s eyes). From these local reconstructions, we find that the distribution of accessible flashes is peaked at short distance, but is also long-tailed and extends much further in certain directions (Fig. S4). Terrain and vegetation also create large variations in the number and range of accessible interactions depending on the firefly’s orientation.

In conclusion, our results have shown that although *P. carolinus* males synchronize locally their rhythmic flashing with their peers, a global swarm synchronization is only possible if enough fireflies are active to transport the collective pace information. Firefly local interactions, rather than metric or topological in nature, are possibly supported by a dynamic network of visual connections defined by relative orientations and visual occlusion from terrain and vegetation [25]. This results in a mixture of short-range and long-range interactions. Such self-organization allows for the possibility for an individual to position itself to be more or less connected, for example by flying above the swarm to be more visible and carry flashing information further. This, in turn, might enable social differentiation. Indeed, it seems *a priori* paradoxical that a group of males competing for female attention exhibits such strong mimicry. While convincing explanations for the ecological function of synchronous flashing have been proposed [26, 27] it is possible to assume that males would use subtle variations in their behavior to distinguish themselves. In particular, further analysis should attempt to understand why and how flash bursts originate at specific locations.

## Acknowledgments

Field experiments were authorized by the National Park Service, permit #GRSM-2020-SCI-2075. We are very grateful to Great Smoky Mountains National Park, notably Becky Nichols and Paul Super, for allowing and facilitating our research. We would like to warmly thank Lynn Faust and Andrew Moiseff for precious guidance and insightful conversations, and Marine de Marcken for commenting on the manuscript. This research was partly funded by the BioFrontiers Institute at CU Boulder.

## Supplementary Materials for

### Materials and Methods

#### Experimental sites

Field experiments took place in the Elkmont, TN area of Great Smoky Mountain National Park, after approval by the National Park Service (permit #GRSM-2020-SCI-2075). In accordance with Park policies, the exact location of the experimental sites will not be disclosed publicly but may be indicated upon request. The general area for field experiments encompasses over 1 mile of trails through densely forested areas, and video recordings took place at several different locations with various camera setups, including in particular the “ridge area”. Extreme care was taken to preserve the local vegetation and wildlife.

#### Three-dimensional reconstruction of the ridge area

For stereoscopic recordings of the ridge area, we used two Sony α7R4 cameras, with settings: 60 frames-per-second (fps); exposure time 1/60 s; maximum aperture (f/1.8); maximum ISO (32,000); and focus at infinity. The cameras were positioned about 4m apart, and their relative orientation was adjusted to optimize the overlap of their fields-of-view (FOV). Using landscape markers such as trees we made sure that each camera’s FOV was similar for each night. Spatial calibration was performed using 5-10 pictures of a checkerboard (25mm square side length) placed at different locations and using the MATLAB stereo calibration toolbox. Temporal calibration (frame synchronization) was based on both a short artificial light signal at the beginning of the recording, and cross-correlation of the temporal patterns between both cameras, which returned the same results. After extraction of the flash positions in each frame using intensity thresholding, triangulation was performed using the MATLAB *triangulate* function to compute the 3D coordinates of recorded flashes.

About 60% of the flashes recorded in each camera could be triangulated. Some mild postprocessing was then applied to eliminate about 1% of outliers points, for example those falling at a distance much greater than 30m from the cameras or at a negative elevation.

#### Three-dimensional reconstruction using 360-degree cameras

The details of the principle and implementation of stereo-vision using 360-degree cameras are described in Ref. (6). Briefly, two GoPro Fusion 360-degree cameras recording at 30fps were placed on the ground at a set distance (0.9m or 1.8m) facing the same direction. The trajectory of a small light was used for calibration, and triangulation was performed using the algorithm provided in Ref. (6). The cameras were started at around 9:30pm each night at various locations across the experimental area and recorded for about 100min.

### Supplementary Text

#### Collective periodicity at high density

Individual *Photinus carolinus* males typically exhibit a flashing pattern consisting of a train of 1 to 6 flashes, each lasting about 0.1s and separated by a 0.55s interflash interval. At high density, swarms of *Photinus carolinus* fireflies produce synchronous flashes that follow a doubly-periodic process of synchronous flashes over the course of ∼10s bursts. While this pattern is clearly visible in the time series of the number of flashes per frame (Fig. 1), the regularity of these two processes is better demonstrated by looking at the corresponding frequency power spectra, *i*.*e*. the Fourier transforms of the time series.

In Fig. S1, we present the power spectra corresponding to 1hr of data for June 3 (low density) and June 11 (high density). At low density, only one frequency peak is present, at about 1.8Hz, corresponding to the typical 0.55s interval between flashes produced by one individual during a flash train. At high density, the same frequency is present, but an emergent “swarm frequency” at about 0.08Hz appears. This denotes the regularity of the flash bursts, which are repeated approximately every 12s. (This value is not the time between the end of a burst and the beginning of the next one, but rather between the maxima of successive bursts.)

Depending on environmental conditions, such as temperature, humidity, hour of the night and progress within the mating season, some slight shifts in these two frequencies can appear.

#### Flashes and “specks”

For increased clarity and accuracy in what follows, we now establish a distinction between “flashes”, which we define as the light emitted by a firefly for about 0.1s, and “specks”, which we define as the impression of a flash on the camera’s sensor over the course of a single frame. Depending on brightness and other factors, a flash consists of a set of up to 8-10 specks, localized in space and on the frame. Due to the finite temporal resolution of videorecording (1/60 s), a speck is associated with a unique time t, equivalent to the number of the frame in which it appears. A flash may span a range of times.

#### Definition of the phase *ϕ*

In order to calculate the relative timing of a speck within a burst, which we denote *ϕ*, we consider the time series of the number *N* of flashes recorded in each frame. Because the time series is noisy, and the beginning of the bursts (with few fireflies) is ill-defined, we rather define the origin of a burst at its center. To identify the time-origin of a burst *b*, we programmatically find the time 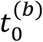 where *N* reaches its maximum within *b*. The phase of a speck at time *t* within the same burst is then simply 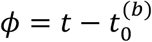. For simplicity, we limit the range of *ϕ* to ±5s.

Using a different method to calculate burst origins, such as a parabolic fit to the spike envelope, does not significantly alter subsequent results.

#### Propagation data for all peak nights

To obtain the propagation plot in Fig. 2D, we proceed as follows. We start from the set of three-dimensional speck coordinates, 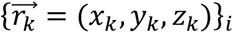, with their associated phases *ϕ*_*k*_ and burst numbers *b*_*k*_. Then, for each pair (*i, j*) of coordinates with the same *b* number and *ϕ*_*i*_< *ϕ*_*j*_, we calculate the Euclidean distance 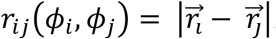.

Subsequently, we group the set of distances *r*_*ij*_into bins in the *ϕ*_*i*_× *ϕ*_*j*_ space. We use bins of side-length 0.5s and discard the diagonal bins for which *ϕ*_*i*_ ≈ *ϕ*_*j*_ because they contain a large fraction of self-correlations (distances between specks corresponding to the same flash). In other word, we group together *r*_*ij*_ that are such that, for instance, (*ϕ*_*i*_, *ϕ*_*j*_)∈ [−5, −4.5[× [−4.5, −4 [or [−5, −4.5[× [−4, −3.5[, …, [−4, −3.5[× [1, −1.5[, etc.

The empirical distributions of *r*_*ij*_ in each bin are displayed in Fig. S3. They all show broad distributions and a mode that increases with the difference between *ϕ*_*j*_ and *ϕ*_*i*_. In Fig. 2D only the distribution median is reported for each bin.

#### Firefly dynamics

To characterize the motion of fireflies, we define streaks as the spatial localization of a flash as it appears on successive frames. From the set of 3D speck coordinates, it is straightforward to concatenate them into distinct realizations of single flashes using a distance-based matching algorithm, since fireflies are rather slow (up to about ∼1m/s, or ∼1cm/frame) and dispersed, hence making matching ambiguities very unlikely.

To calculate average streak velocities, we first remove all streaks consisting of 4 or less speck positions. For the remaining streaks, we simply calculate the velocity vector by subtracting the first coordinate to the last one, and rescaling to get m/s units. Fig. 3B is obtained by calculating the empirical distributions of streak velocities as a function of their corresponding average phase.

In Fig. S2, we use the distribution of streak velocities over the propagation event shown in Fig. 2B,C to show that although flash information propagates along the *y*-axis, there is no flow of fireflies along that same axis.

#### Immersive 3D reconstruction

To estimate line-of-sight interactions from the perspective of a swarming firefly, *i*.*e*. seen from a point of view close to the ground and surrounded by vegetation, we use stereoscopic setups of pair of 360-degree cameras. 360-degree cameras record in every direction around them, and hence can be placed directly within the swarm, instead of outside of it.

In Fig. S4, we present the immersive 3D reconstructions from two different sites (FS & TC) with different surrounding environments. The spatial distributions of accessible flashes show that the number and spatial range of interactions vary significantly depending on the orientation of the probe firefly. Local interactions are peaked at short distance, but also extend significantly to large distances. Due to visual occlusion, some directions allow to see a large number of peers, while others are much more obstructed. Directions that are more open also increase the range of accessible interactions. In the presence of a terrain elevation, the average distance of accessible flashes increases significantly.

**Fig S1.**
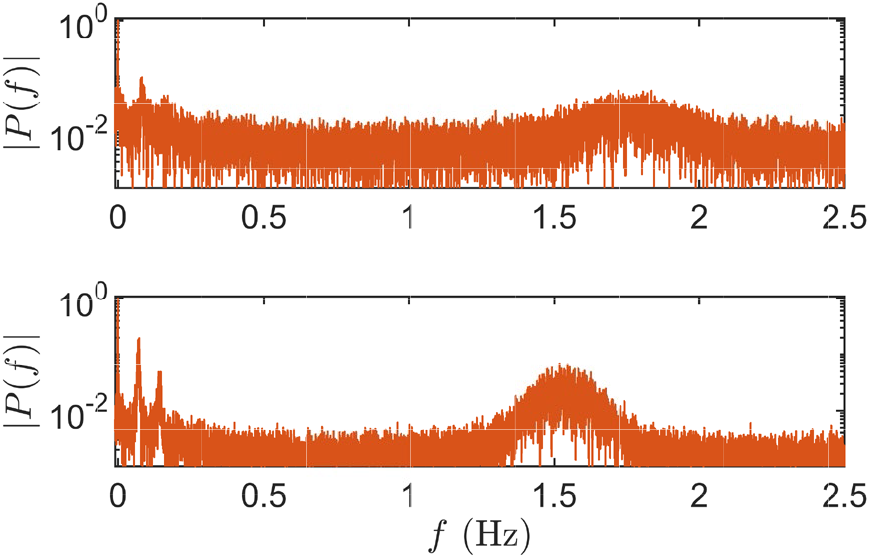
Fourier Transforms of the *N*-time series for June 3 (top) and June 11 (bottom) (1hr of recording between 22:00 and 23:00 EST). The frequency peak between 1.5-2Hz corresponds to the time between successive flashes in a flash train. At high density (June 11), an emergent low frequency appears around 0.08Hz, corresponding to burst periodicity.

**Fig S2.**
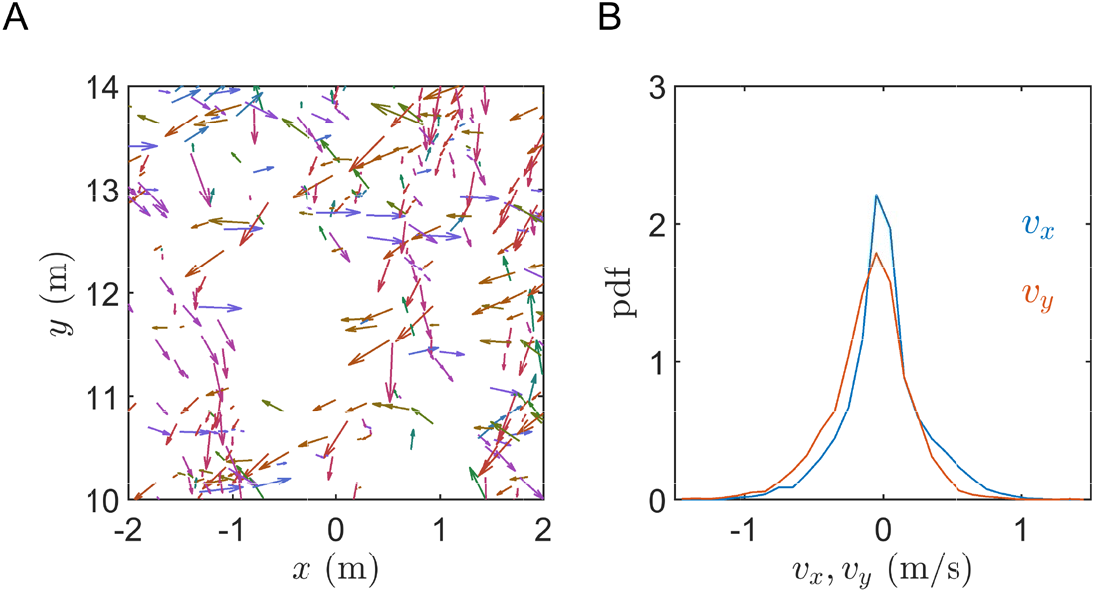
Streak velocities for the propagation event shown in Fig. 2B,C. **(A)** Sample of streak locations and velocities (direction and magnitude) in the horizontal plane. No preferential directionality appears towards the +*y* axis, which is the direction of propagation of the burst wave. **(B)** Distribution of velocities along the *x*- and y- axes (about 3000 vectors). Both distributions are narrow and centered. The mean ⟨*v*_*y*_⟩ ≈ −0.1 m/s, which is in the opposite direction of flash propagation, and small compared to the corresponding standard deviation, ≈0.3 m/s. Therefore, there is no evidence of a significant flow of fireflies parallel to flash propagation.

**Fig S3.**
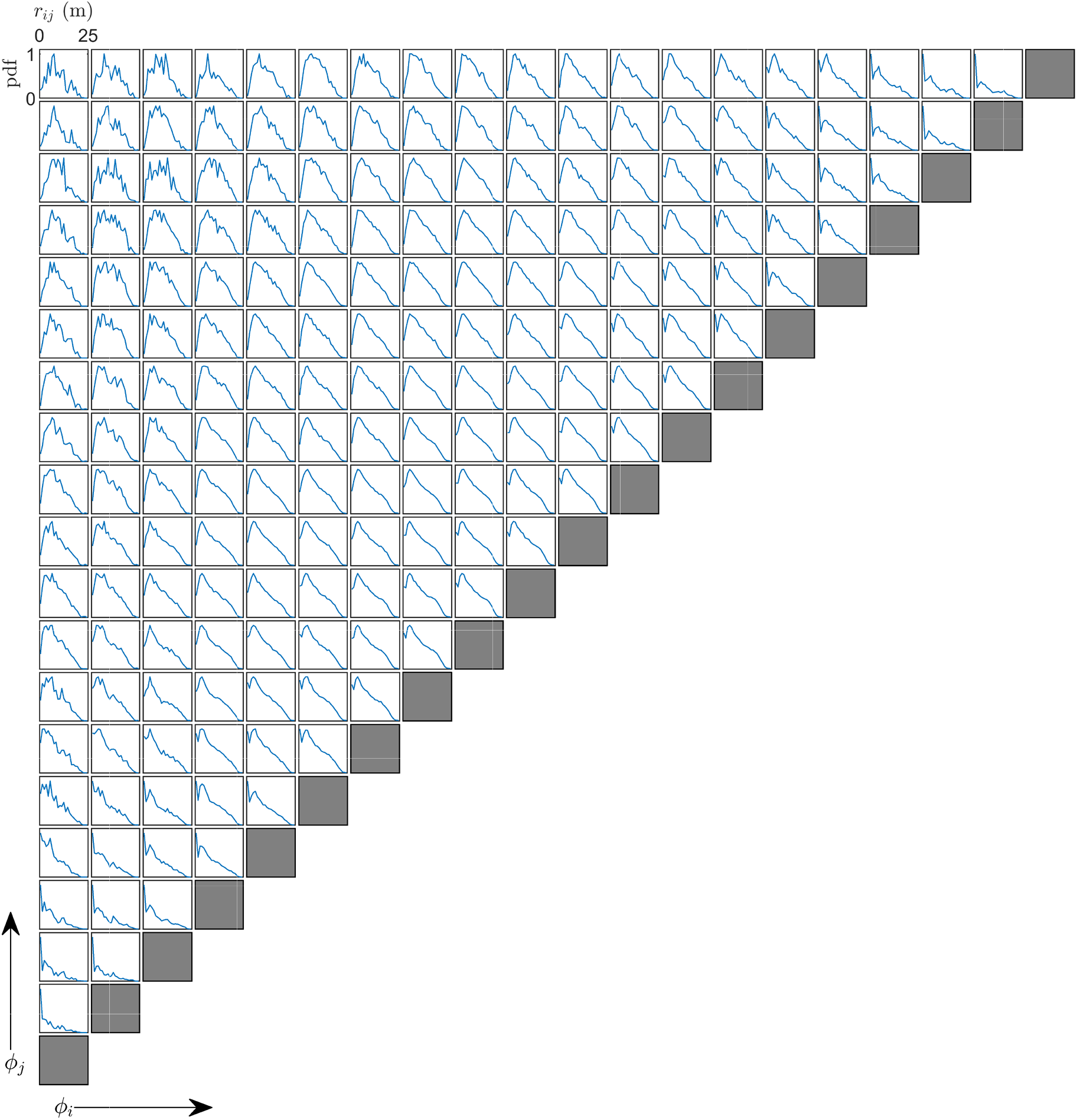
Empirical probability density functions (pdf, normalized by their maximum value for consistent scale) for the flash distances *r*_*ij*_ (*ϕ*_*i*_, *ϕ*_*j*_) in the *ϕ*_*i*_ × *ϕ*_*j*_ space discretized into bins of (0.5s)^2^. The *ϕ* intervals are the same as in Fig. 2D, which displays the median values of each of the above distributions. Data is shown for full recordings (about 2hr30min) on June 11.

**Fig S4.**
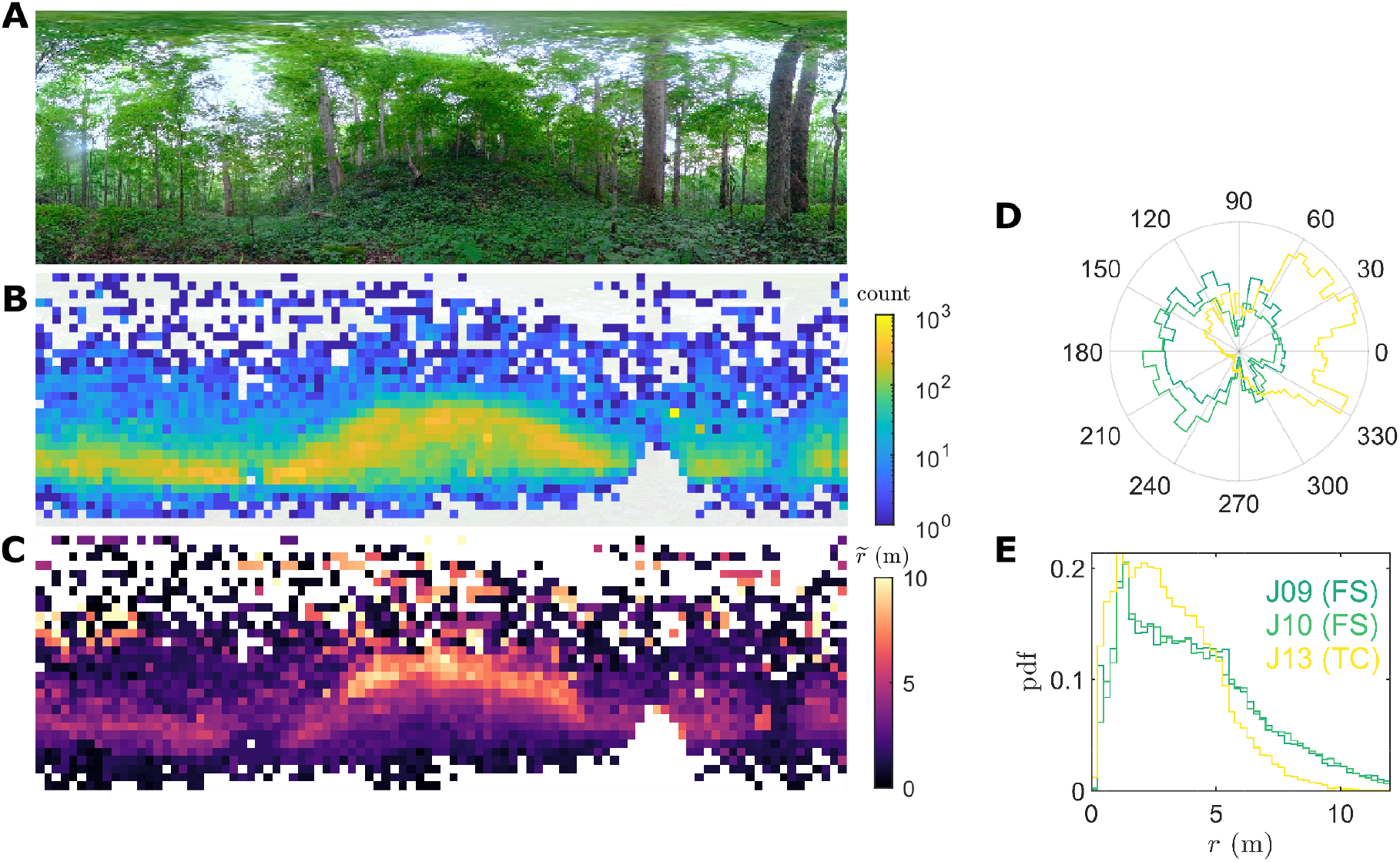
Illustration of local interactions from the perspective of a swarming firefly, reconstructed from 360-degree stereo-recordings (90min). **(A)** Stereographic projection of the view in one 360-degree camera, showing the terrain and obstruction from trees and vegetation (site TF, June 10). **(B)** Corresponding spatial distribution of flashes. Fireflies are localized primarily in the horizontal plane. Fewer flashes are accessible along visually-occluded directions, *e*.*g*. where the FOV is obstructed by trees or bushes. **(C)** Median distance 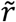 of accessible flashes as a function of direction. Flashes that occur at an elevated position, either on a hill or flying above ground, can typically be seen from further away. In obstructed areas, perceived flashes remain close. This is how both short-range and long-range visual interactions can co-exist. **(D)** Distribution of flashes as a function of angle for 3 different sets of recordings (2 at FS, 1 at site TC). Visual occlusion creates strong disparities in the number of accessible flashes, depending on the orientation of the firefly’s FOV. **(E)** Distribution of flash distances for the same recordings. Due to terrain heterogeneity and occlusion, most flashes are concentrated at short distance, but a large fraction extend much farther.

**Movie S1**.

This movie shows a 1min segment of the propagation event described in Fig. 2B,C. Reconstructed flash locations are displayed in real time in the horizontal plane. Colors indicate the phase, and the progressing time series at the top shows the corresponding timing within the burst. Early flashes occur generally at small y, and as the burst progresses, flashes move towards the increasing y, *i*.*e*. far from the cameras and towards the top of the ridge.

**Movie S2**.

This movie shows all the flash occurrences captured in the 10min segment displayed in Fig. 2B,C, as a function of their burst phase *ϕ* and projected in the horizontal plane. Because the spatiotemporal pattern in Movie S1 is noisy (the time-distance relationship is broad), it is fruitful to make use of the burst periodicity to “average out” multiple propagation paths. The average burst temporal pattern is displayed at the top.

